# Machine learning guided design of high affinity ACE2 decoys for SARS-CoV-2 neutralization

**DOI:** 10.1101/2021.12.22.473902

**Authors:** Matthew C. Chan, Kui. K. Chan, Erik Procko, Diwakar Shukla

## Abstract

A potential therapeutic candidate for neutralizing SARS-CoV-2 infection is engineering high-affinity soluble ACE2 decoy proteins to compete for binding of the viral spike (S) protein. Previously, a deep mutational scan of ACE2 was performed and has led to the identification of a triple mutant ACE2 variant, named ACE2_2_.v.2.4, that exhibits nanomolar affinity binding to the RBD domain of S. Using a recently developed transfer learning algorithm, TLmutation, we sought to identified other ACE2 variants, namely double mutants, that may exhibit similar binding affinity with decreased mutational load. Upon training a TLmutation model on the effects of single mutations, we identified several ACE2 double mutants that bind to RBD with tighter affinity as compared to the wild type, most notably, L79V;N90D that binds RBD with similar affinity to ACE2_2_.v.2.4. The successful experimental validation of the double mutants demonstrated the use transfer and supervised learning approaches for engineering protein-protein interactions and identifying high affinity ACE2 peptides for targeting SARS-CoV-2.

---

Severe acute respiratory syndrome coronavirus-2 (SARS-CoV-2) is a positive-sense RNA-enveloped virus that causes coronavirus disease 2019 (COVID-19) and has since emerged as a global pandemic^1^. As of December 2021, the COVID-19 outbreak has resulted in over 250 million reported cases and claimed over 5 million deaths around the world^2^. Even with the successful development of mRNA-based vaccines, issues regarding vaccine hesitancy, vaccine availability in low- and middle-income countries, and emerging variants of concern presents a dire need for alternative therapeutic options for COVID-19.

Extensive work has characterized the primary mechanism of SARS-CoV-2 entry to be facilitated by the interactions involving the spike (S) protein and the angiotensin converting enzyme 2 (ACE2) receptor^3,4^. S is trimeric class I viral fusion glycoprotein that consists of soluble S1 and membrane-tethered S2 subunits. The S1 subunit contains the receptor-binding domain (RBD) which, when directly binds with the peptidase domain of ACE2, initiates the cleavage and shedding of S1, thereby exposing the S2 subunit for membrane fusion and viral RNA release into the host cell^5^. As such, S is a major target for anti-SARS-CoV-2 therapeutics, including monoclonal antibodies that compete with ACE2 for RBD binding and neutralize virus infection^5^. However, mutations in S present in emerging variants of SARS-CoV-2, including B.1.1.7 (Alpha), B.1.351 (Beta), P.1 (Gamma), G.1.617.2 (Delta), and BA.1 (Omicron) enables SARS-CoV-2 to bind to ACE2 with greater affinity than the wild-type strain^6,7^. The N501Y mutation identified in the Alpha, Beta, Gamma, and Omicron strains introduces additional π-π and cation-π interactions with ACE2 residues that increase the binding affinity by 10-fold as compared to the wild-type RBD^8^. Furthermore, the E484K mutation in the Beta and Gamma variants introduces new electrostatic interactions with neighboring acidic residues on ACE2^9^. Consequently, due to these altered molecular interactions, SARS-CoV-2 variants are more likely to evade the immune response and reduce the efficacy of neutralizing antibodies and vaccines^10^.

Efforts in developing therapeutic alternatives have focused on engineering soluble ACE2 (sACE2) constructs that acts as a high-affinity decoy for the viral S protein. sACE2, in which the transmembrane domain is removed, has been previously shown to be capable of binding to the S protein and is currently undergoing clinical trails for treating COVID-19^3,11^. Recently, Chan and colleagues performed a deep mutational scan of ACE2 and uncovered several amino acid substitutions throughout the ACE2 receptor that enhance the binding affinity to S of SARS-CoV-2^12^. They further characterized a sACE2 derivative containing the three mutations T27Y, L79T, and N330Y, referred to as sACE2_2_.v2.4 (**Figure 1A**), to exhibit a 35-fold increase in binding affinity to SARS-CoV-2, and further tighter affinity for SARS-CoV-2 variants that contain the N501Y mutation^12,13^. *In vivo* characterization further demonstrates that inoculation of sACE2_2_.v2.4 in mice prevents acute lung vascular endothelial injury induced by SARS-CoV-2 and improves survival^14^. While the mutations introduced in sACE2_2_.v2.4 were chosen intuitively, it presents an opportunity to identify other potential high-affinity sACE2 derivatives, specifically double mutants, with a decreased mutational load.

**Figure 1:**
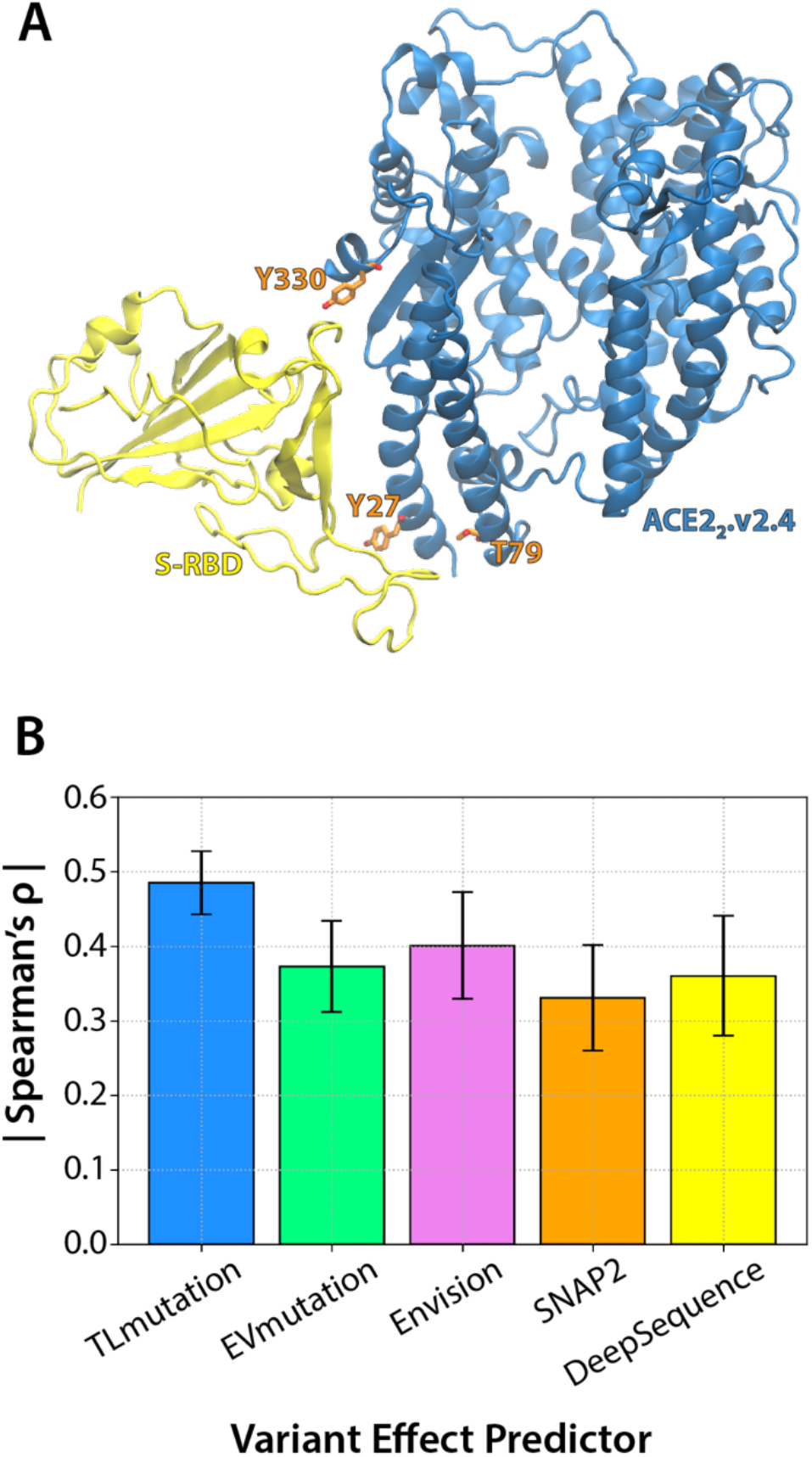
(A) Model of ACE2_2_.v.2.4-RBD complex based on the resolved ACE2-RBD structure (PDB: 6M17). ACE2 is shown in blue cartoon representation. RBD domain is shown as yellow cartoon. Introduced mutations (T27Y, L79T, N330Y) in ACE2_2_.v.2.4 are shown in orange sticks. Figure was generated using VMD 1.9.3^23^ (B) Performance of variant effector predictors in predicting ACE2 single mutants for RBD binding. Spearman’s rank correlation (ρ) is measured as the average correlation across a 20% test or withheld dataset, across five folds.

Given that the search space of ACE2 double mutants consist of over two million combinations, it is impractical to experimental evaluate all possible mutants. Therefore, we employed an *in silico* screening using our previous described algorithm TLmutation^15^ to identify ACE2 double mutant derivatives that might achieve tight binding to the SARS-CoV-2 S protein. The TLmutation platform is an adaptation of the variant effect predictor EVmutation (part of the EVCoupling package) which uses sequence and evolutionary information to determine the effect of a mutation based on a residue’s site-specific conservation and coevolutionary coupled interactions^16^. However, as the effect of a mutation predicted by EVmutation is a measure of overall survivability, the native function of the protein may be masked and may not necessarily correlate to a specific function of study. To overcome this, the TLmutation platform implements a supervised algorithm to train a model with the experimental scores from a deep mutational scan. In doing so, the weights of the site-specific conservation and coevolutionary couplings are decoupled from a function of overall survival and fitted to the experimentally tested function of the protein. We have previously demonstrated the use of TLmutation to successfully incorporate deep mutational scanning data to decouple protein function from overall survival to enhance the prediction of variant effects in 13 individual proteins^15^. More applicable to this current study, the TLmutation algorithm supersedes the performance of the EVmutation for predicting the effect of higher order mutations^15^.

We first constructed a EVmutation model trained on a multiple sequence alignment consists of sequences above 80% sequence identify and totaled in over 8,000 sequences. As we expect this model to gauge protein survival rather than binding function, we then construct a TLmutation model by training the preliminary model on the 2,340 single point mutation obtained from deep mutagenesis of the human ACE2 receptor scanned for enhanced binding of the RBD domain^12^. The performance of the single mutation model was evaluated using a 5-fold cross validation in which the dataset was randomly split into an 80/20 train/test set. Across five folds, the experimental effect of ACE2 single mutations on RBD binding had the highest correlation with the TLmutation score compared to other variant effect predictors^16–19^ (**Figure 1B**). As such, the TLmutation algorithm demonstrates the ability to decouple overall survivability and predict a particular function of interest. We find that DeepSequence, which models the evolutionary information into a latent representation^19^ and is considered a top preforming method for predicting deep mutational scans^20^, performed subpar to TLMutation (**Figure 1B**). We suspect that as ACE2 binding to S is not evolutionary favored, the multiple sequence alignment used to train DeepSequence and EVmutation does not discriminate mutations that may optimize RBD binding.

As TLmutation utilizes coevolutionary couplings to make predictions, we used the model trained on single point mutations to predict the effect of ACE2 double mutants. To validate the predictions generated by the model, we selected 18 ACE2 double mutants from the following different categories: 9 double mutants that scored in the top 1% of TLmutation; 4 double mutants in which each single mutation independently enhances binding, but scored less than the top 1%; 2 double mutants with a TLmutation score in the lowest 1% to serve as negative controls; and 3 double mutants consisting of the combination of single mutations in sACE2_2_.v2.4 (**Table S1**). The ACE2 mutants were evaluated by expression of full-length, myc-tagged protein in human Expi293F cells, which were tested by flow cytometry for binding to superfolder GFP (sfGFP)-fused RBD (**Figure 2**). The predicted constructs exhibited levels of expression similar with the wild-type ACE2, suggesting these mutants do not significantly alter the stability of the receptor (**Figure S1**). With the exception of the two negative controls, all the mutants tested were observed to bind better or on par with the wild type (**Figure 2**). Additionally, the TLmutation scores correlated more to the experimental results as compared to EVmutation and DeepSequence (TLmutation R^2^ = 0.52, EVmutation R^2^ = 0.09, DeepSequence R^2^ = 0.23, **Figure 3A-C**).

**Figure 2:**
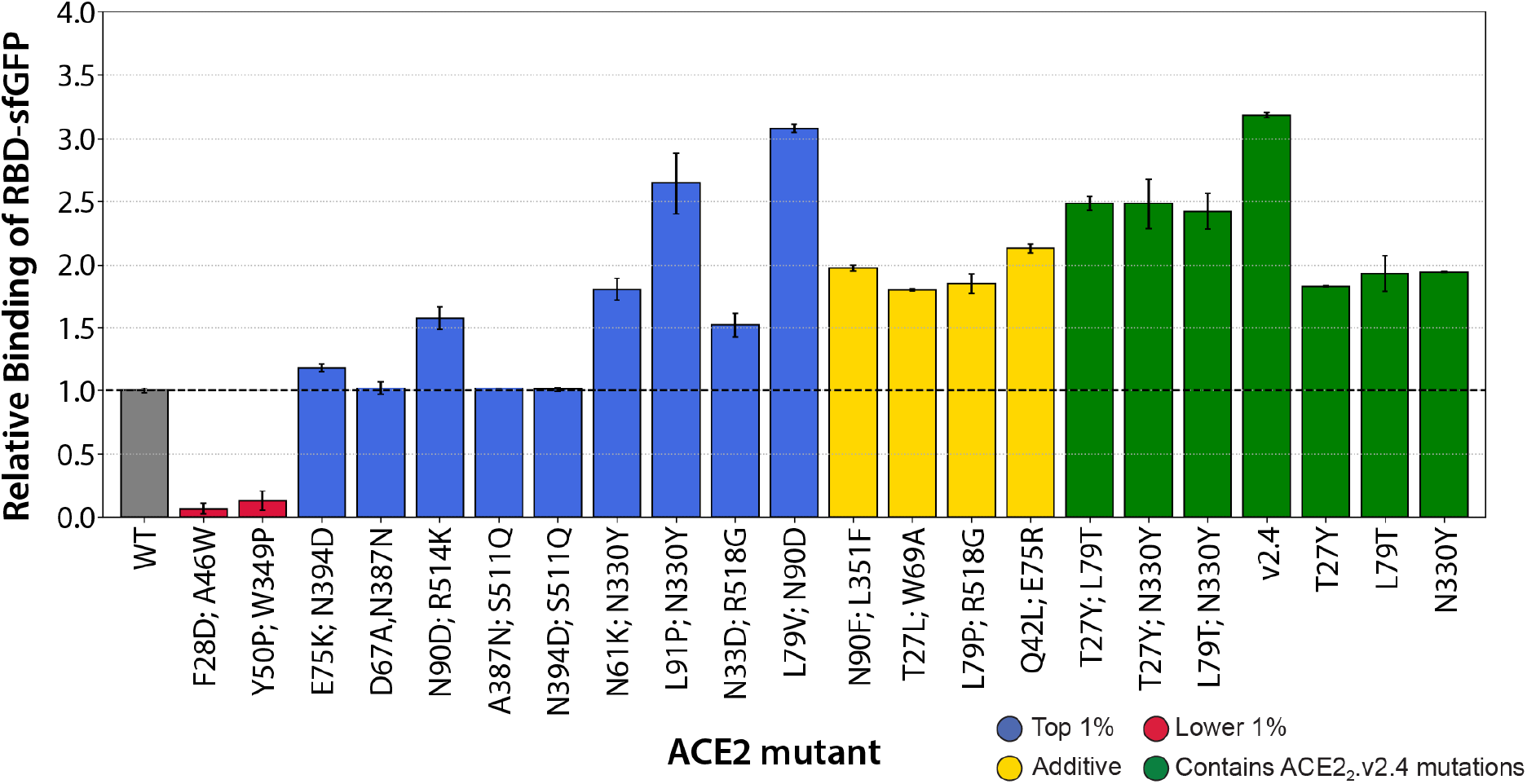
Performance of ACE2 double mutants predicted by TLmutation. Expi293F cells expressing wild type or mutant myc-ACE2 were co-incubated with a fluorescent anti-myc antibody and with RBD-sfGFP expression medium at a sub-saturating dilution. After gating for a constant level of anti-myc fluorescence to control for expression differences, bound RBD-sfGFP was measured by flow cytometry. Data are mean, N=2, error bars represent range. The double mutants belonged to 4 groups and are colored accordingly: blue, mutants that scored in the top 1% of TLmutation; red, that scored in the lowest 1% of TLmutation; yellow, double mutants whose single mutations were previously validated to significantly enhance binding; green, double mutants of the mutations in ACE2_2_.v2.4.

**Figure 3:**
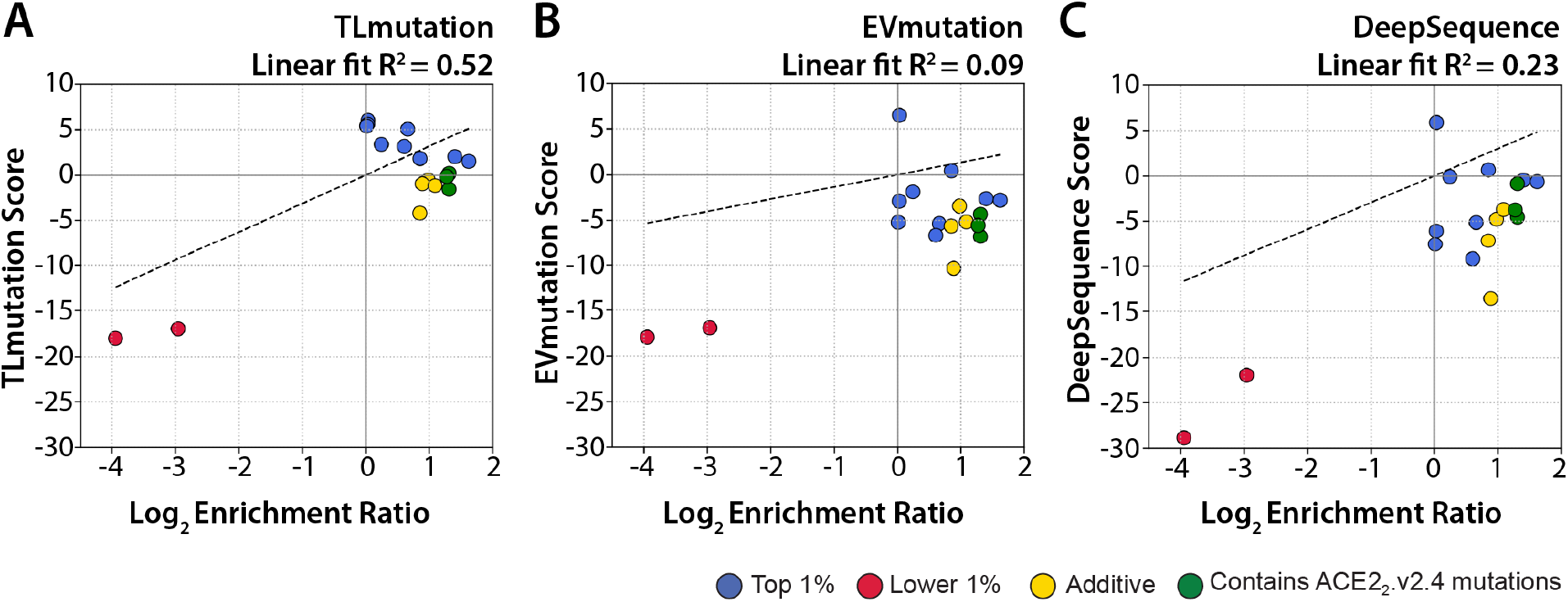
Comparison of variant effect predictions TLmutation (B), EVmutaion (C) and DeepSequence (D) with experimental measurements. Points are colors according to the 4 groups of double mutants and are colored accordingly: blue, mutants that scored in the top 1% of TLmutation; red, that scored in the lowest 1% of TLmutation; yellow, double mutants whose single mutations were previously validated to significantly enhance binding; green, double mutants of the mutations in ACE2_2_.v2.4..

**Figure 4:**
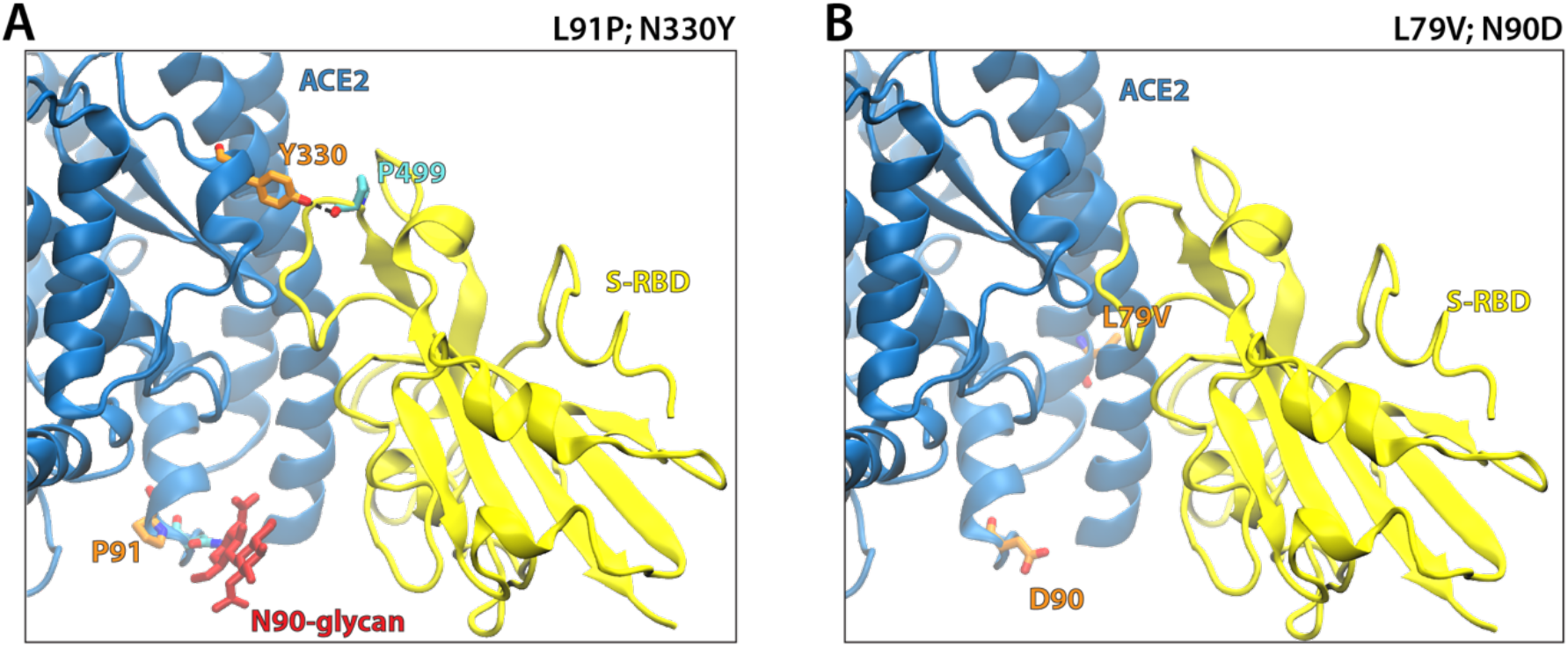
Structural models of the top two ACE2 double mutants of high RBD binding. (A) The L9IP;N330Y variant. Y330 may interact with P499 of RBD, while P91 may enable helical kinks to allosterically stabilize the interfacial helices. Alternatively, P91 may also alter the dynamics of the N90 glycan, which has been shown to shield RBD binding to ACE2^21^. (B) The L79V;N90D variant. D90 disrupts the glycan which would promote RBD binding.

Double mutants containing the combination of sACE2_2_.v2.4 mutations, T27Y;L79T, T27Y;N330Y, and L79T;N330Y, bound to RBD with greater affinity than the single mutant variants, and had similar affinity as sACE2_2_.v2.4. When examining the *in silico* mutagenesis predictions for these double mutants, TLmutation score the sACE2_2_.v2.4 double mutants in the top 14%, with T27Y;N330Y and L79T;N330Y scoring in the upper 5%. DeepSequence scored all three mutants in the top 11%, whereas in EVmutation, it was score in the top 25% of all double mutants. When incorporating the newly experimental results of double mutants to the original single mutation training set, TLmutation, EVmutation, and DeepSequence predicted sACE2_2_.v2.4 to be highly deleterious in function. We hypothesize the limited dataset of double mutations was not adequate to distinguish critical couplings to accurately predict higher order mutations.

For the nine mutants that scored in the top 1% of the TLmutation predictions, three mutants D67A;N387N, A387N;S511Q, and N394D;S511Q failed to be distinguishable from the wild type. The remaining mutants from the same group were observed to perform better than the wild type. Two ACE2 mutants exhibited over a two-fold relative binding increase over the wild type. For the L91P;N330Y mutant, both single mutations exhibited a relative binding fold enhancement of under two^12^; however, when expressed together, the relative binding increased over 2.5 fold. We have previously conducted molecular dynamics simulations of sACE2_2_.v2.4 with the RBD domain of SARS-CoV-2 and observed the N330Y mutation to interact with backbone carbonyls of P499 on RBD further to stabilize the associated complex^14^ (**Figure 4A**). Interestingly, L91P is not directly located on the ACE2 binding interface and may allosterically stabilize the α2 helix for favorable S binding^12^. It is also possible that the proline introduction may alter the dynamics of the glycosylation motif at position 90. The best performing double mutant, L79V;N90D (**Figure 4B)**, bound RDB-sfGFP exceptionally well, equivalent to sACE2_2_.v2.4. This double mutant scored within the top 0.7% of all TLmutation double mutants, whereas in EVmutation and DeepSequence, it scored in the top 5^th^ and 2^nd^ percentile, respectively. However, this mutant disrupts the N90 glycan which has been shown to interact with RBD^21^ and could impair glycosylation that is required for *in vivo* compatibility, solubility, and reduced drug immunogenicity in ways that are unpredictable. Overall, the TLmutation algorithm successfully predicted higher order ACE2 mutants with tight RBD binding activity from single mutation data, but nonetheless the sACE2_2_.v2.4 derivative remains a top candidate for drug development as it combines high affinity spike protein binding and preserves the native glycosylation sites.

With regards to engineering a soluble ACE2 peptide for high RBD binding activity, the search space of possible combination of mutations is unfathomable to experimentally evaluate. Furthermore, identifying variants of high activity cannot be solely obtained from intuition, sequence conservation, or single mutational data alone (**Figure S2**). Indeed, many high scoring (above a TLmutation score of 2 or the top 0.4% of all scored variants) ACE2 double mutants contains residues not involved at the interface (**Figure S3**). As such, computational techniques that employ *in silico* mutagenesis provide a powerful and attractive approach to systematically identify potential candidates for further validation. A recent *in silico* mutagenesis study employing the Rosetta based modeler for protein-protein association, Flex ddG, has successfully identified a quadruple ACE2 mutant, S19F;T27F;K31W;N330F, to exhibit nanomolar binding affinity similar to the sACE2_2_.v2.4 designed mutant^22^. Similar in this study, the use of a supervised transfer learning algorithm implemented by TLmutation successfully identify several double mutations that enhances binding to the RBD domain of SARS-CoV-2. Though, while the mutations initially identified sACE2_2_.v2.4 construct retains the very tight binding activity and provide the most *in vivo* compatibility, these works demonstrate the successful use of computational techniques to aid in the engineering of protein design. A current limitation of variant effect predictors, TLmutation included, is in its ability to predict higher order mutations due to the lack of training data to identify relevant residue couplings. As mentioned, TLmutation, EVmutation, and DeepSequence fail to predict sACE2_2_.v2.4 variant to be functional in binding. However, as deep mutational datasets become more readily available, specifically double mutational scans, we expect the use of computational algorithms may not only improve *in silico* variant effect predictions, but further aid in the systematic design of protein function that considers *in vivo* compatibility with minimal mutational load.

## Methods

### In silico mutagenesis

We performed an *in silico* screening of all possible ACE2 double mutants using our previous described machine learning algorithm TLmutation. The TLmutation algorithm utilizes the EVmutation platform in addition to deep mutational scanning data of single mutations to predict the ACE2 mutational landscape for RBD binding. The TLmutation algorithm was first trained on the experimental single mutations data^12^ to decouple the weights from overall organismal survivability of fitness. The weights were then optimized to fit the function under investigation, i.e. RBD binding. The performance of the algorithm was evaluated with a 5-fold cross validation in which the data were split into 80/20 train/test sets. The TLmutation predictions were compared to predictions from the EVcouplings package^16^, Envision web server^17^, the SNAP2 web server^18^, and the DeepSequence package^19^. For construction of the EVmutation, TLmutation, and DeepSequence model, the multiple sequence alignment was constructed using the EVcouplings package. TLmutation and EVmutation was performed using Python 3.7 on CentosOS6.6 with access to AMD EPYC 7301 (1200 MHz, 126 GB RAM). DeepSequence was performed using Python 2.7 on Ubuntu 20.04.3 with access to an Nvidia GeForce RTX 2080TI (11GB RAM) and an Intel i9-10900K (3700 MHz, 32 GB RAM). The input human ACE2 sequence was truncated to contain only positions 10 to 529 to reduce computational cost.

### Assessment of ACE2 mutations from TLmutation

Expi293F cells transfected with pCEP4-myc-ACE2 plasmids were collected 24 h posttransfection (600 × g, 60 s), washed with ice-cold Dulbecco’s phosphate buffered saline (PBS) containing 0.2% bovine serum albumin (BSA), and stained with 1:50 RBD-sfGFP expression medium (prepared as previously described and 1:250 anti-myc Alexa 647 (clone 9B11, Cell Signaling Technology) in PBS-BSA. After 30 minutes incubation on ice, cells were washed twice in PBS-BSA and analyzed on a BD Accuri C6 using instrument software. Gates were drawn around the main population based on forward and side scatter properties, followed by a gate for medium Alexa 647 fluorescence to control for any expression differences between mutants. Mean superfolder GFP (sfGFP) fluorescence was measured and background fluorescence of negative cells was subtracted. All replicates were from independent, distinct samples.

## Supporting information

Supplementary Results

## Acknowledgements

This work is supported by the C3.ai Digital Transformation Institute Research Award provided by C3.ai Inc and Microsoft Corporation, NIH grant R35GM142745, and intramural funds of the University of Illinois College of Medicine to D.S, and NIH grants R43-AI162329 to E.P. and K.K.C.

