## Supplementary Results for "Machine learning guided design of high affinity ACE2 decoys for SARS-CoV-2 neutralization"

### Supplemental Figures

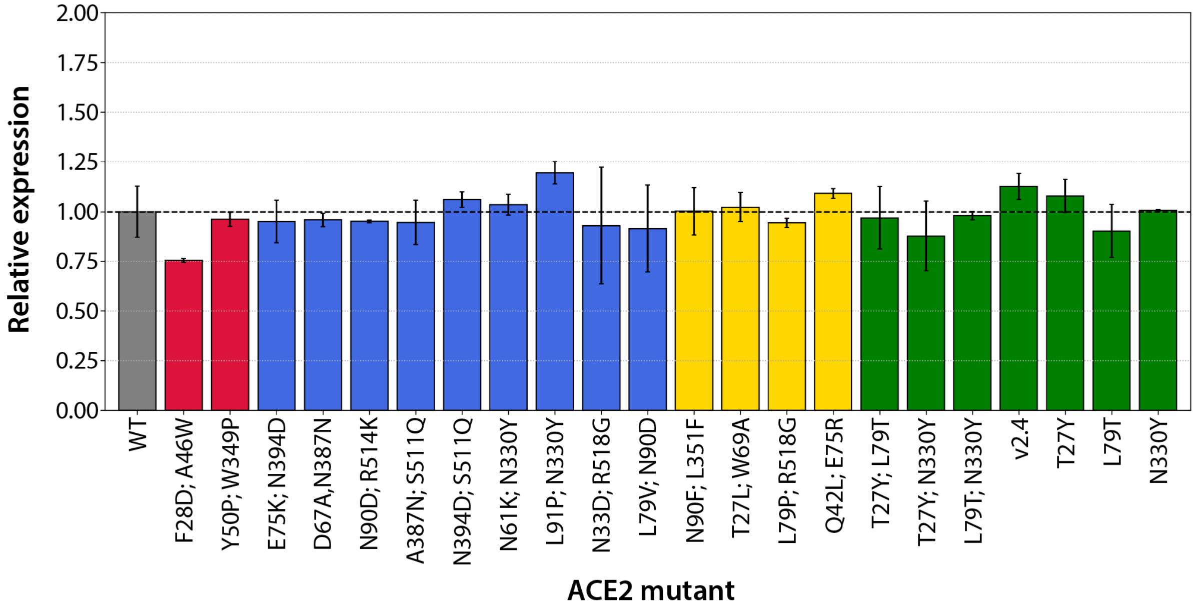


**Figure S1:** ACE2 expression measure by detection of the extracellular myc tag. Data are mean, N=2, error bars represent range. The double mutants belonged to 4 groups and are colored accordingly: blue, mutants that scored in the top 1% of TLmutation; red, that scored in the lowest 1% of TLmutation; yellow, double mutants whose single mutations were previously validated to significantly enhance binding; green, double mutants of the mutations in $ACE2_2$.v2.4.

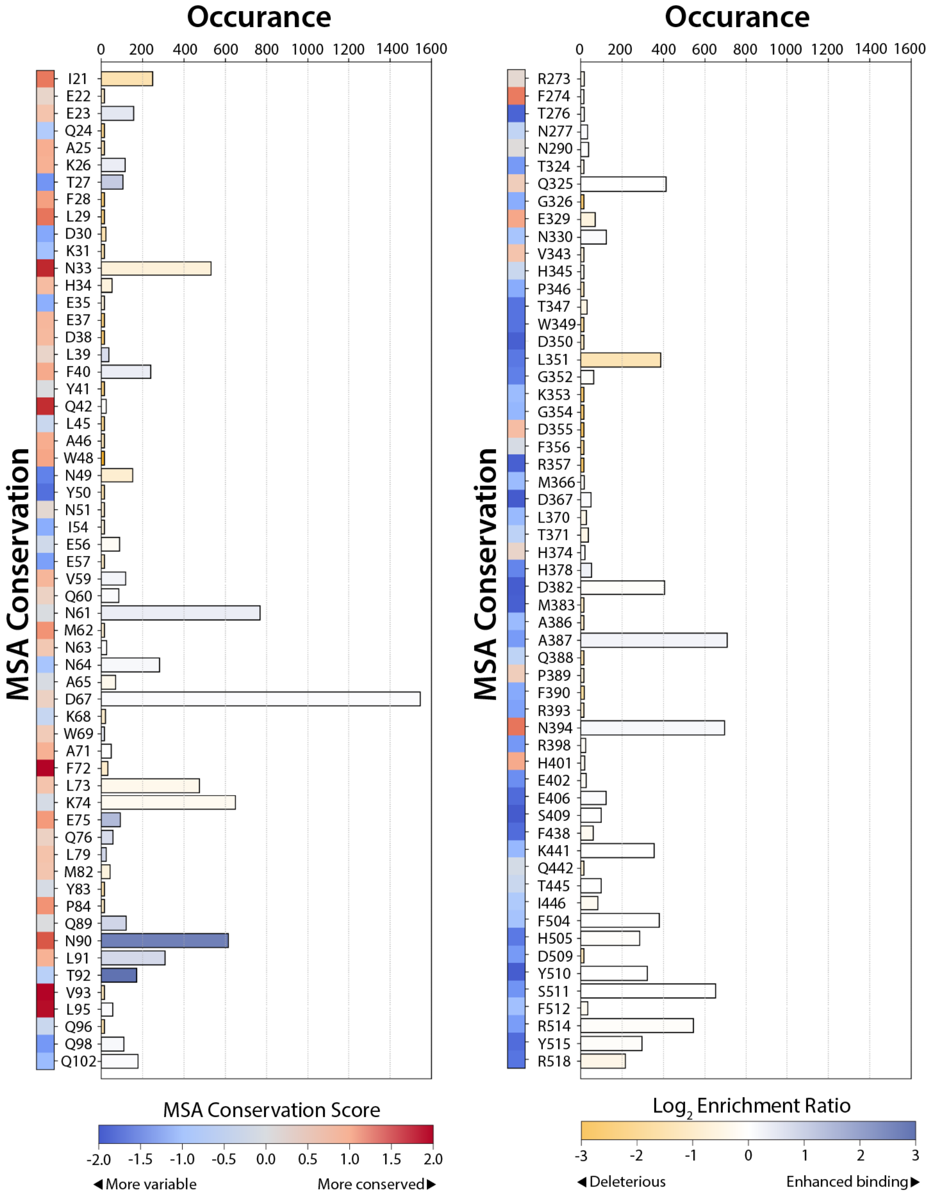


**Figure S2:** Occurrence of ACE2 residues positions that scored above the TLmutation score of 2 or the top 0.4% of predicted ACE2 double mutants. The top 0.4% of ACE2 double mutants consisted of 7,563 ACE2 variants. The occurrence a particular residue is involved in the 7,563 ACE2 variance is represented in bars, with the bar color depicting the average $\log_2$ enrichment ratio of ACE2-RBD binding from the deep mutational scan and colored from deleterious (orange) to enhanced activity (blue). The conservation of the residue at each position is shown to the left of the residue label and colored from variable (blue) to conserved (red). Multiple sequence alignment (MSA) conservation score was generated using the ConSurf database.

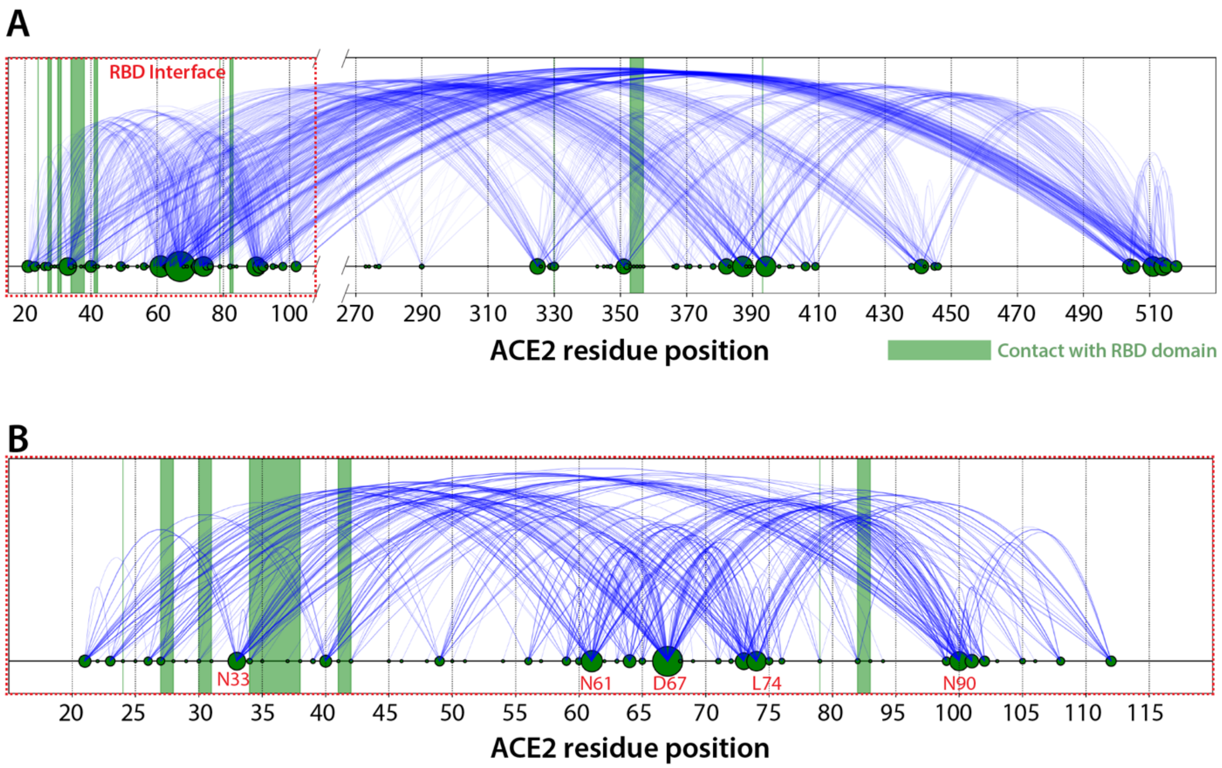


**Figure S3:** Frequency of ACE2 residues pairs that scored above the TLmutation score of 2 or the top 0.4% of predicted ACE2 double mutants. The top 0.4% of ACE2 double mutants consisted of 7,563 ACE2 variants. (A) The frequency of residue pairs in 7,563 ACE2 double mutants is represented as blue lines and involves residues beyond the ACE2-RBD binding interface (red dashed square). Domains of direct contact with RBD highlighted in green rectangles. Relative frequency of residue position is represented by circle size. (B) Zoomed-in representation of the frequency of residue pairs involving the first three helices that form the ACE2-RBD interface.

**Supplemental Table**

Table S1: Comparison of *in silico* mutagenesis predictions. Scores and percentile of the 18 double mutants experimental tested for sfGFP-RBD binding are reported for the variant effect predictors TLmutation, EVmutation, and DeepSequence (DeepSeq). Color indicates 4 groups of double mutants tested: red, that scored in the lowest 1% of TLmutation; blue, mutants that scored in the top 1% of TLmutation; yellow, double mutants whose single mutations were previously validated to significantly enhance binding; green, double mutants of the mutations in $ACE2_2$.v2.4.

| Mutant | Relative binding w.r.t WT | TLmutation score | TL percent | EVmutation score | EV percent | DeepSeq score | DeepSeq percent |
|---|---|---|---|---|---|---|---|
| WT | 1.000 | N/A | N/A | N/A | N/A | N/A | N/A |
| F28D; A46W | 0.065 | -18.077 | 99.97% | -17.960 | 99.57% | -28.864 | 98.63% |
| Y50P; W349P | 0.129 | -17.017 | 99.85% | -16.943 | 98.68% | -21.973 | 89.28% |
| E75K; N394D | 1.184 | 3.340 | 0.05% | -1.900 | 2.86% | -0.101 | 1.35% |
| D67A; A387N | 1.022 | 5.997 | < 0.01% | 6.490 | < 0.01% | 5.860 | 0.01% |
| N90D; R514K | 1.580 | 5.033 | < 0.01% | -5.388 | 14.98% | -5.155 | 12.43% |
| A387N; S511Q | 1.020 | 5.587 | < 0.01% | -2.931 | 5.00% | -6.104 | 16.14% |
| N394D; S511Q | 1.008 | 5.383 | < 0.01% | -5.241 | 14.16% | -7.617 | 22.85% |
| N61K; N330K | 1.805 | 1.792 | 0.52% | 0.438 | 0.41% | 0.643 | 0.83% |
| L91P; N330Y | 2.640 | 2.002 | 0.40% | -2.661 | 4.32% | -0.473 | 1.69% |
| N33D; R518G | 1.522 | 3.117 | 0.07% | -6.678 | 23.85% | -9.235 | 30.83% |
| L79V; N90D | 3.078 | 1.491 | 0.75% | -2.829 | 4.72% | -0.643 | 1.87% |
| N90D; L351F | 1.975 | -0.552 | 6.60% | -3.465 | 6.53% | -4.795 | 11.14% |
| T27L; W69A | 1.802 | -4.217 | 40.43% | -5.643 | 16.50% | -7.201 | 20.93% |
| L79P; R518G | 1.852 | -0.963 | 9.15% | -10.384 | 57.98% | -13.595 | 54.31% |
| Q42L; E75R | 2.128 | -1.168 | 10.62% | -5.198 | 13.90% | -3.720 | 7.77% |
| T27Y; L79T | 2.481 | -1.568 | 13.85% | -4.363 | 9.86% | -0.869 | 2.13% |
| T27Y; N330Y | 2.482 | 0.175 | 3.15% | -6.813 | 24.92% | -4.560 | 10.36% |
| L79T; N330Y | 2.419 | -0.175 | 4.67% | -5.580 | 16.13% | -3.767 | 7.90% |
| v2.4 | 3.182 | -0.629 | N/A | -8.378 | N/A | -26.648 | N/A |
| T27Y | 1.832 | N/A | N/A | N/A | N/A | N/A | N/A |
| L79T | 1.932 | N/A | N/A | N/A | N/A | N/A | N/A |
| N330Y | 1.944 | N/A | N/A | N/A | N/A | N/A | N/A |
